# LincRNA-Cox2 functions to regulate inflammation in alveolar macrophages during acute lung injury

**DOI:** 10.1101/2021.07.15.452529

**Authors:** Elektra Kantzari Robinson, Atesh K. Worthington, Donna M. Poscablo, Barbara Shapleigh, Mays Mohammed Salih, Haley Halasz, Lucas Seninge, Benny Mosqueira, Valeriya Smaliy, E. Camilla Forsberg, Susan Carpenter

**Affiliations:** Department of Molecular, Cell and Developmental Biology, University of California Santa Cruz, 1156 High St, Santa Cruz, CA 95064; Institute for the Biology of Stem Cells, University of California-Santa Cruz, Santa Cruz, California, United States of America; Department of Biomolecular Engineering, University of California Santa Cruz, 1156 High St, Santa Cruz, CA 95064

## Abstract

The respiratory system exists at the interface between our body and the surrounding non-sterile environment; therefore, it is critical for a state of homeostasis to be maintained through a balance of pro- and anti- inflammatory cues. An appropriate inflammatory response is vital for combating pathogens, while an excessive or uncontrolled inflammatory response can lead to the development of chronic diseases. Recent studies show that actively transcribed noncoding regions of the genome are emerging as key regulators of biological processes, including inflammation. LincRNA-Cox2 is one such example of an inflammatory inducible long noncoding RNA functioning to control immune response genes. Here using bulk and single cell RNA-seq, in addition to florescence activated cell sorting, we show that lincRNA-Cox2 is most highly expressed in the lung, particularly in alveolar macrophages where it functions to control immune gene expression following acute lung injury. Utilizing a newly generated lincRNA-Cox2 transgenic overexpressing mouse, we show that it can function in trans to control genes including Ccl3, 4 and 5. This work greatly expands our understanding of the role for lincRNA-Cox2 in host defense and sets in place a new layer of regulation in RNA-immune-regulation of genes within the lung.

## Introduction

Acute lung injury (ALI) and it’s more severe form, known as acute respiratory distress syndrome (ARDS), are caused by dysregulated inflammatory responses resulting from conditions such as sepsis and trauma (Moldoveanu *et al*, 2009; Mokra & Kosutova, 2015; Wang *et al*, 2019b; Mowery *et al*, 2020; Butt *et al*, 2016). Fundamentally, the characteristics of ALI include neutrophilic alveolitis, dysfunction of barrier properties, microvascular thrombosis, the formation of hyaline membrane, alveolar macrophage dysfunction, as well as indirect systemic inflammatory responses (Pittet *et al*, 1997; Alluri *et al*, 2017; Gouda & Bhandary, 2019; Fan & Fan, 2018). Although a variety of anti-inflammatory pharmacotherapy are available, the morbidity and outcome of ALI/ARDS patients remain poor (Raghavendran *et al*, 2008; Yin & Bai, 2018; Suo *et al*, 2018; Deng *et al*, 2017). Therefore, obtaining a more complete understanding of the molecular mechanisms that drive ALI inflammatory dysfunction is of great importance to improving both the diagnosis and treatment of the condition.

Long noncoding RNAs (lncRNAs) are a class of non-coding RNAs that include 18,000 in human and nearly 14,000 in the mouse genome (Uszczynska-Ratajczak *et al*, 2018; Fang *et al*, 2018). Since their discovery, lncRNAs have been shown to be key regulators of inflammation both *in vitro* and *in vivo* (Robinson *et al*, 2020; Statello *et al*, 2021). Moreover, lncRNAs have also been characterized to be stable and detectable in body fluids (Quinn & Chang, 2016), and therefore have enormous potential for biomarker discovery in both diagnosis and prognosis applications (Aftabi *et al*, 2021; Ma *et al*, 2021; Viereck & Thum, 2017). A number of studies have been carried out to better understand the gene regulatory network between lncRNAs and mRNAs during ALI to identify novel biomarkers (Teng *et al*, 2021; Wang *et al*, 2019a). In addition to searching for biomarkers for ALI there have been studies performed to try and understand the functional mechanisms for lncRNAs during ALI (Chen *et al*, 2021). Lipopolysaccharide (LPS)-induced acute lung injury (ALI) is a commonly utilized animal model of ALI as it mimics the inflammatory induction and polymorphonuclear (PMN) cell infiltration observed during clinical ALI (Asti *et al*, 2000). One study showed that knocking down MALAT1, a well-studied lncRNA, exerts a protective role in the LPS induced ALI rat model and inhibited LPS-induced inflammatory response in murine alveolar epithelial cells and murine alveolar macrophages cells through sponging miR-146a (Dai *et al*, 2018). Additionally, Xist has been shown to attenuate LPS-induced ALI by functioning as a sponge of miR-146a-5p to mediate STAT3 signaling (Li *et al*, 2021).

Previous work by ourselves and others identifies long intergenic noncoding RNA-Cox2 (lincRNA-Cox2) as a regulator of immune cell signaling in macrophages (Carpenter *et al*, 2013; Tong *et al*, 2016; Hu *et al*, 2016; Covarrubias *et al*, 2017; Hu *et al*, 2018; Liao *et al*, 2020; Xue *et al*, 2019). We have previously characterized multiple mouse models to show that lincRNA-Cox2 functions *in vivo* to regulate the immune response. We showed that lincRNA-Cox2 knockout mice (Sauvageau *et al*, 2013) have profound defects in the neighboring protein coding gene Ptgs2. We went on to show that lincRNA-Cox2 regulates Ptgs2 in *cis* through an enhancer RNA mechanism requiring locus specific transcription of the lncRNA (Elling *et al*, 2018). In order to study the function of lincRNA-Cox2 independently of its role in regulating Ptgs2 in *cis* we generated a lincRNA-Cox2 mutant mouse using CRISPR to target the splice sites resulting in significant loss of the RNA allowing us to study the role for the RNA in *trans*. We performed an LPS-induced endotoxic shock model and confirmed that lincRNA-Cox2 is an important positive and negative regulator of immune genes in *trans*. We previously showed that lincRNA-Cox2 is most highly expressed at steady-state in the lung and in this study, we utilize our mutant model to determine if lincRNA-Cox2 can function in trans to regulate gene expression in the lung (Elling *et al*, 2018). In addition, we characterize a transgenic overexpressing mouse model of lincRNA-Cox2 and show that the defects in immune gene expression caused by removal of lincRNA-Cox2 can be rescued by the transgenic overexpression of lincRNA-Cox2 in an LPS-induced ALI model. Mechanistically, we show through bone marrow chimeric studies that lincRNA-Cox2 expression is coming from bone marrow derived cells to regulate genes within the lung. Finally, we show that lincRNA-Cox2 is most highly expressed in alveolar macrophages where it functions to regulate inflammatory signaling. Collectively we show that lincRNA-Cox2 is a trans-acting lncRNA that functions to regulate immune responses and maintain homeostasis within the lung at baseline and upon LPS-induced ALI.

## Results

### Immune gene expression is altered in the lungs of lincRNA-Cox2 deficient mice

In order to determine the role for lincRNA-Cox2 in the lung we first examined our RNA-sequencing data to compare gene expression profiles in the lungs of WT versus lincRNA-Cox2 mutant mice (Elling *et al*, 2018, GSE117379) (Figure 1A). We found 85 genes down-regulated and 41 genes up-regulated in the lincRNA-Cox2 mutant lungs compared to WT (Figure 1B). Gene-ontology analysis showed that both the down and up regulated genes were associated with the immune system, metabolism, and response to stimulus (Figure 1C-D), which are similar to pathways that lincRNA-Cox2 has previously been associated with in bone marrow derived macrophages (BMDMs) (Carpenter *et al*, 2013). To determine if loss of lincRNA-Cox2 impacts protein expression we performed ELISAs on lung homogenates from WT and lincRNA-Cox2 deficient mice (Figure 1E). While many genes remained unchanged at baseline between the WT and lincRNA-Cox2 mutant lungs (SFigure 1I-P), we did find that Il-12p40, Cxcl10, Ccl3, Ccl4, Cxcl2, Ccl5 and Ccl19 are all significantly up-regulated in the mutant lungs at baseline (Figure 1A-H). Interestingly none of these cytokines are up-regulated at the RNA level in our whole lung tissue RNA-sequencing suggesting that they might be regulated post-transcriptionally or that they are only regulated in a small subset of cells that cannot be easily captured from the whole lung lysate RNA-seq data (SFigure 1I-P).

**Figure 1:**
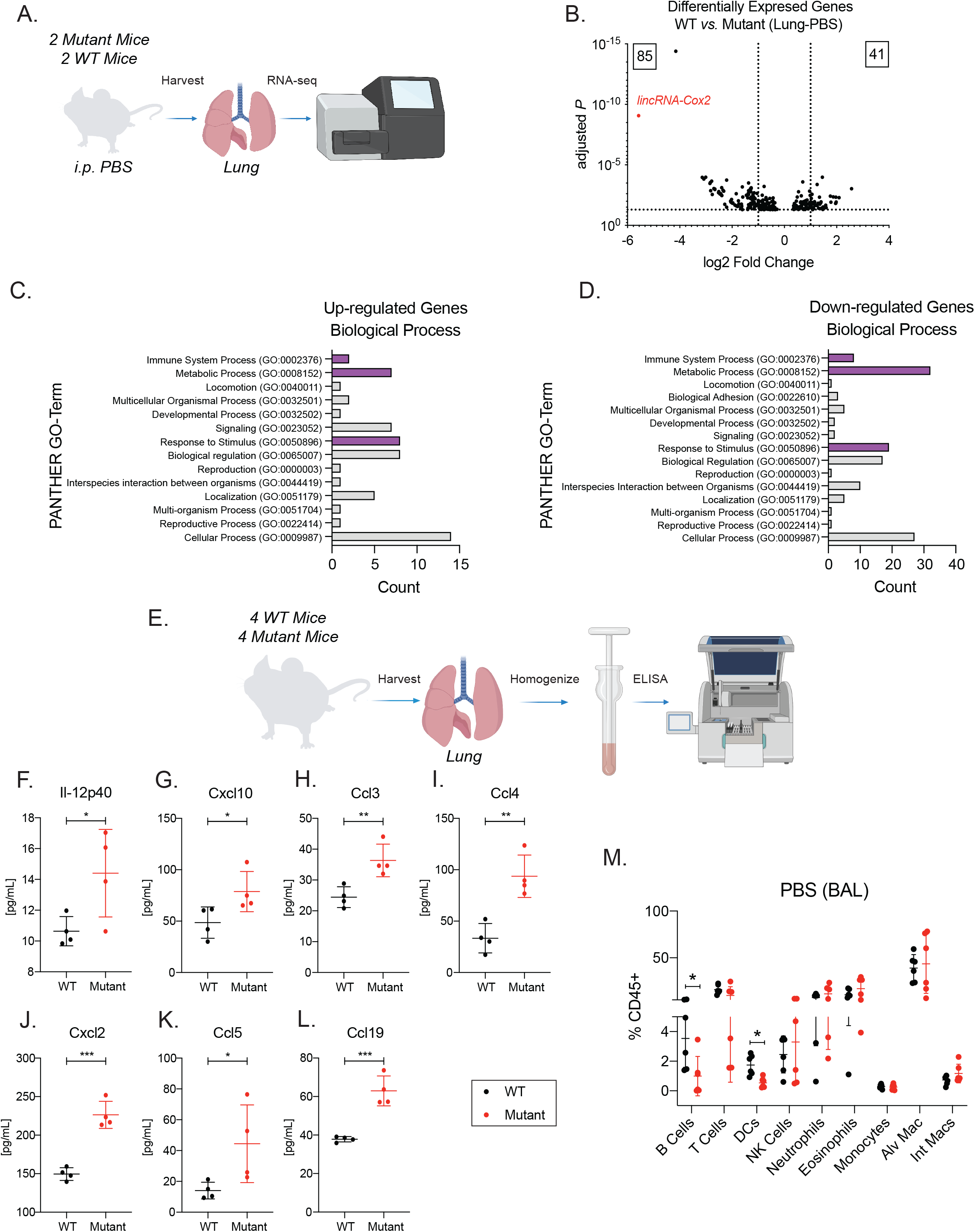
lincRNA-Cox2 regulates immune signaling within the lung during homeostasis. (A) Schematic of RNA-seq analysis of WT and Mutant lungs at baseline. (B) Volcano plot of differentially expressed genes from WT *vs.* Mutant lungs. Biological process gene ontology of (C) up-regulated genes and (D) down-regulated genes. (E) Schematic of cytokine analysis of lung homogenates from WT and mutant mice. Multiplex cytokine analysis was performed on lung homogenates for (F) Il-12p40 (G) Cxcl10 (H) Ccl3 (I) Ccl4 (J) Cxcl2 (K) Ccl5 and (L) Ccl19. (M) Flow cytometry analysis of immune cells in the bronchiolar lavage fluid (BAL) at baseline gated off of CD45+ cells. The student’s t-test used to determine the significance between WT and mutant mice. Asterisks indicate statistical significance (*=> 0.05, **>=.01, ***=> 0.0005).

Finally, we measured the immune cell repertoire in the bronchiolar lavage fluid (BAL) of WT and lincRNA-Cox2 deficient mice and found that B cells and dendritic cells, while at very low expression levels in both strains, are significantly lower at baseline in the mutant mice (Figure 1M). These findings indicate that lincRNA-Cox2 functions as both a positive and negative regulator of immune gene expression which can impact the cellular milieu within the lung at steady state.

### LincRNA-Cox2 regulates the pro-inflammatory response during acute lung injury (ALI)

Our data suggest that lincRNA-Cox2 plays a role in regulating immune gene expression at steady-state, therefore we wanted to determine if loss of lincRNA-Cox2 could impact the immune response during acute lung injury. We employed an LPS induced acute lung injury (ALI) model as outlined in Figure 2A. We assessed the immune cell repertoire within the BAL by flow cytometry in WT and lincRNA-Cox2 mutant mice following LPS challenge and found that the most abundant and critical cell type, neutrophils, were significantly reduced when lincRNA-Cox2 is removed (Figure 2B). We also assessed the cytokine and chemokine response both in the BAL and serum of WT and lincRNA-Cox2 mutant mice by ELISA. We found that Il6, Ccl3 and Ccl4 were downregulated in the serum and the BAL of the lincRNA-Cox2 mutant mice (Figure 2C-E). In addition, several proteins were specifically affected either in the serum or BAL, such as Ccl5 and Ccl22 which were upregulated in the serum (Figure 2F-G), while Ifnb1 was upregulated only in the BAL of the mutant mice (Figure 2H). Consistent with our previous work tnf remains unchanged between WT and lincRNA-Cox2 mutant mice (Figure 2I). These data suggest that lincRNA-Cox2 exerts different effects within the lung compared to the periphery and that lincRNA-Cox2 can impact acute inflammation at the protein and cellular levels within the lung.

**Figure 2:**
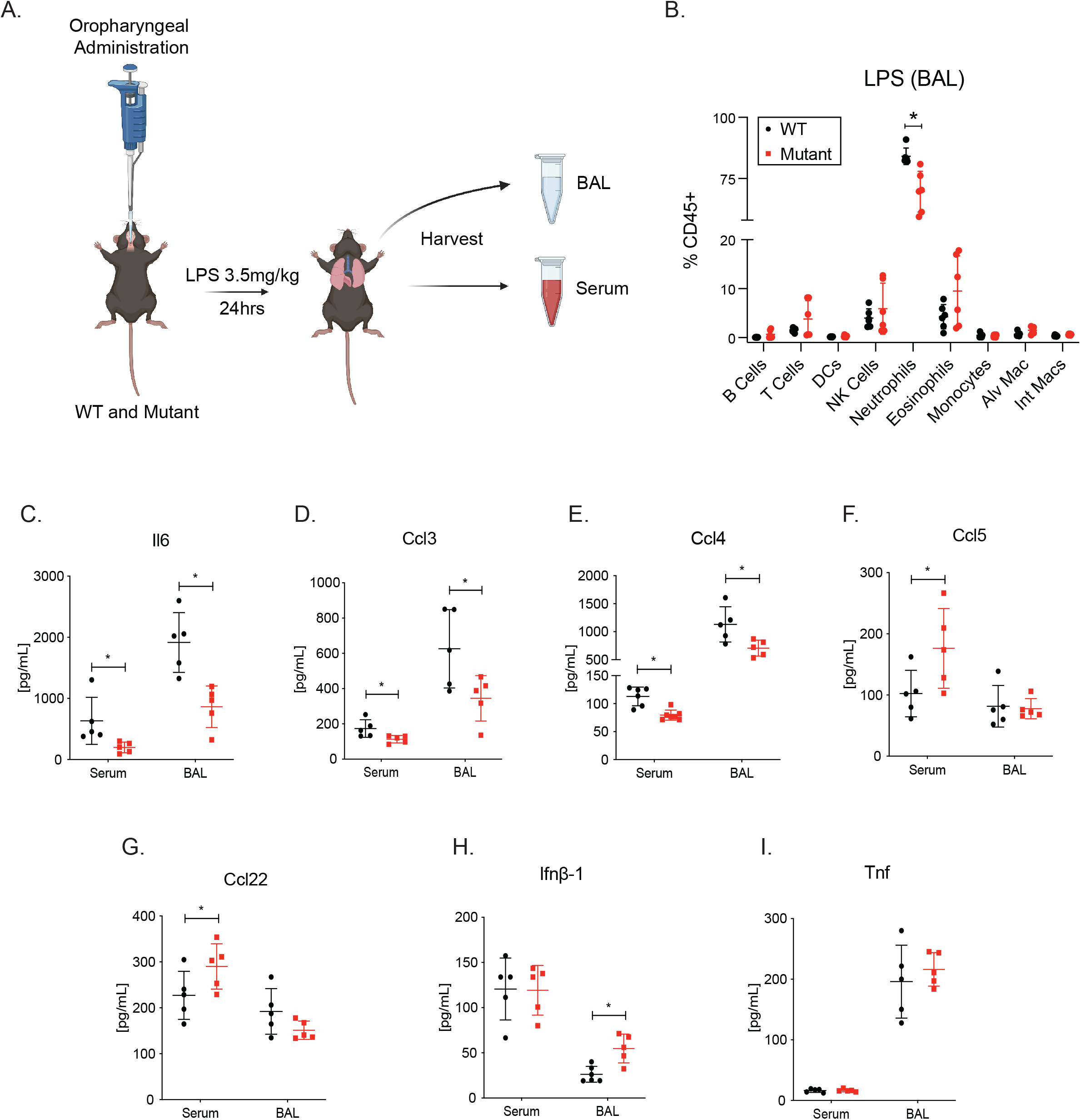
lincRNA-Cox2 positively regulates the pro-inflammatory response during acute lung injury (ALI). (A) ALI schematic depicting the oropharyngeal route of 3.5mg/kg LPS administration in WT and Mutant. Mice were sacrificed after 24 h, followed by harvesting serum and bronchiolar lavage fluid (BAL). (B) Body temperatures were measured at 0 h, 6 h, 12 h, and 24 h. (C) BAL cells were analyzed by flow cytometry to assess recruitment of immune cells in WT and immune cells gated off CD45+. Multiplex cytokine analysis was performed on serum and BAL for (D) Il6, (E) Ccl5, (F) Ccl3, (G) Ccl4, (H) Ccl22, (I) Ifnb-1 and (J) Tnf. The student’s t-test used to determine the significance between WT and mutant mice. Asterisks indicate statistical significance (*=> 0.05, **>=.01, ***=> 0.0005).

### Generation and characterization of a transgenic mouse model overexpressing lincRNA-Cox2

We have determined that lincRNA-Cox2 is critical in regulating inflammation at baseline, during a septic shock model (Elling *et al*, 2018) and here during an acute lung injury model. In order to confirm that lincRNA-Cox2 is functioning in *trans*, we generated a transgenic lincRNA-Cox2 mouse line using the TARGATT system, which allows for stable integration of lincRNA-Cox2 into the H11 locus (Tasic *et al*, 2011) (Figure 3A). The inserted cassette is carrying a CAG promoter, lincRNA-Cox2, and an SV40 polA stop cassette. Mice were bred to homozygosity (SFigure 2), and lincRNA-Cox2 levels were measured in WT and Transgenic bone-marrow-derived macrophages (BMDMs) (Figure 3B). As expected, lincRNA-Cox2 is highly expressed in the transgenic macrophages with no difference in expression following LPS stimulation. Next, we performed a septic shock model of WT and transgenic mice to determine if overexpression of lincRNA-Cox2 can impact the immune response (Figure 3C). As expected lincRNA-Cox2 is highly expressed in the lung tissue of the transgenic mice and interestingly we observe increased levels of Il6 while other lincRNA-Cox2 target genes, such as Ccl5 are not affected. This suggests that overexpression of lincRNA-Cox2 can have the opposite phenotype to knocking out the gene to regulate Il6 within the lung (Figure 3D, SFigure 3A-C). We harvested serum from the mice at steady state and found higher levels of Csf1 and lower levels of Il10 (Figure 3E-F) in the mice overexpressing lincRNA-Cox2. Other inflammatory cytokines including Il6, Ccl5, Ccl3 and Ccl4 were unaltered in lincRNA-Cox2 transgenic mice serum (Figure 3G-J). These data suggests that at steady state overexpression of lincRNA-Cox2 does not have broad impacts on gene expression, however it can impact specific genes including Il6 in the lung and Il10 and Csf1 in the periphery.

**Figure 3:**
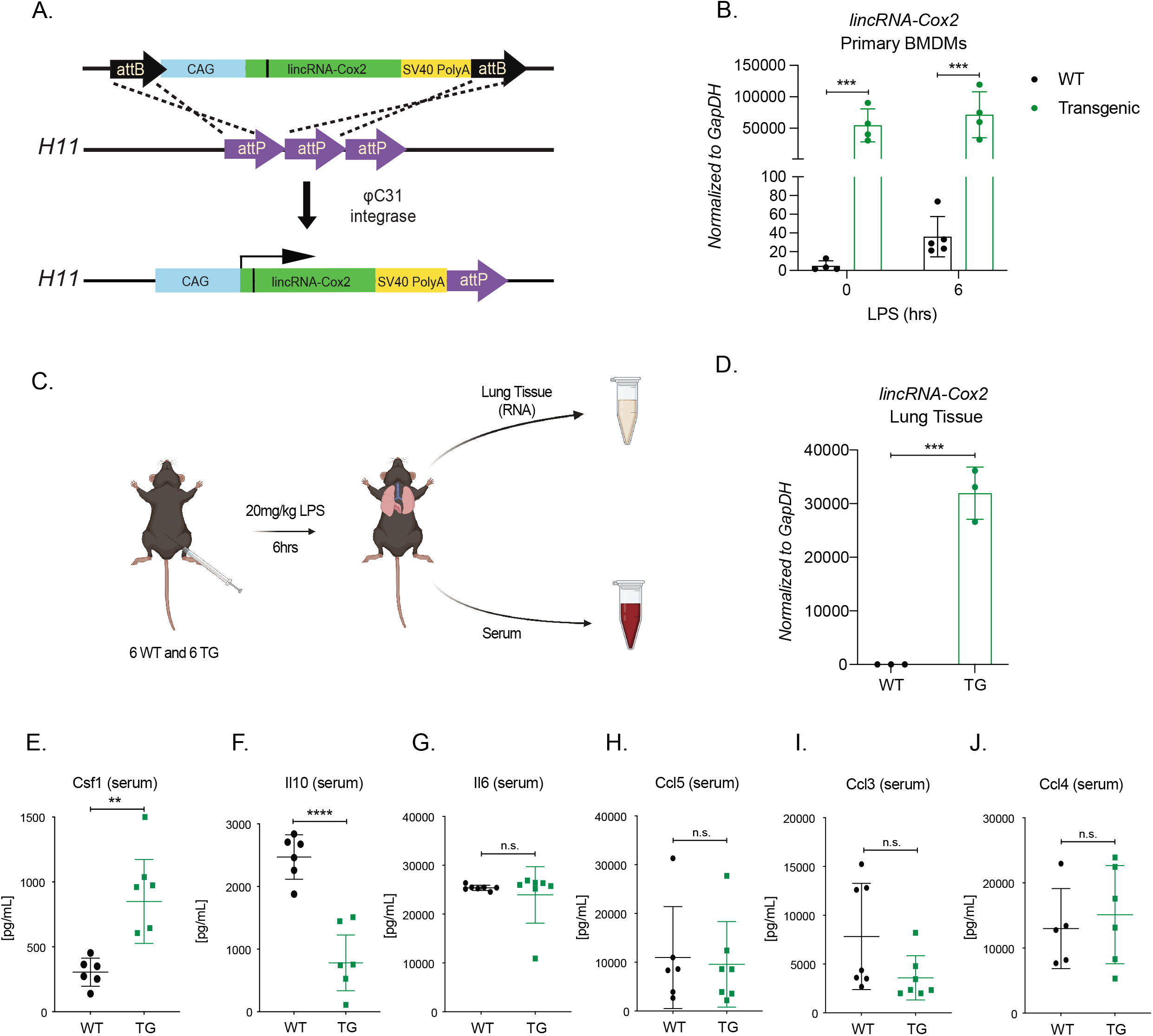
Characterization of lincRNA-Cox2 transgenic mouse. (A) We have generated a transgenic lincRNA-Cox2 mouse line using the TARGATT system. This approach allows for stable integration of lincRNA-Cox2 into the H11 locus. Our inserted cassette is carrying a CAG promoter, lincRNA-Cox2, and an SV40 polA stop cassette. (B) lincRNA-Cox2 levels measured in WT and Transgenic bone-marrow-derived macrophages with and without LPS for 6 h, normalized to GapDH. (C) Schematic of 20mg/kg LPS septic shock model of WT and transgenic mice. (D) lincRNA-Cox2 measured by RT-qPCR in lung tissue. Serum was harvested and ELISAs were performed to measure (E) Csf1, (F) Il10, (G) Il6, (H) Ccl5, (I) Ccl3 and (J) Ccl4. The student’s t-test used to determine the significance between WT and TG mice. Asterisks indicate statistical significance (*=> 0.05, **>=.01, ***=> 0.0005).

### LincRNA-Cox2 functions in trans to regulate acute inflammation

To determine if lincRNA-Cox2 can function in *trans* to regulate immune genes following an *in vivo* challenge with LPS we crossed the mice deficient in lincRNA-Cox2 (Mut) with the transgenic overexpressing mice (TG) generating mice labeled throughout as MutxTG (Figure 4A). We first performed an intraperitoneal (IP) endotoxic shock model to determine if we could rescue the lincRNA-Cox2 phenotype identified in our previous study (Elling *et al*, 2018) (Figure 4B). As expected, lincRNA-Cox2 expression is significantly reduced in the lung tissue and BAL of the deficient mice (Mut) mice, and highly expressed in the MutxTG mouse (Figure 4C-D). We found that Ccl5 and Cxcl10 are expressed at higher levels in the BAL and serum of lincRNA-Cox2 mutant mice (Figure 4E-H) and the expression levels can be rescued by trans expression of lincRNA-Cox2 in the MutxTG mice where the levels of the two proteins return to WT levels.

**Figure 4:**
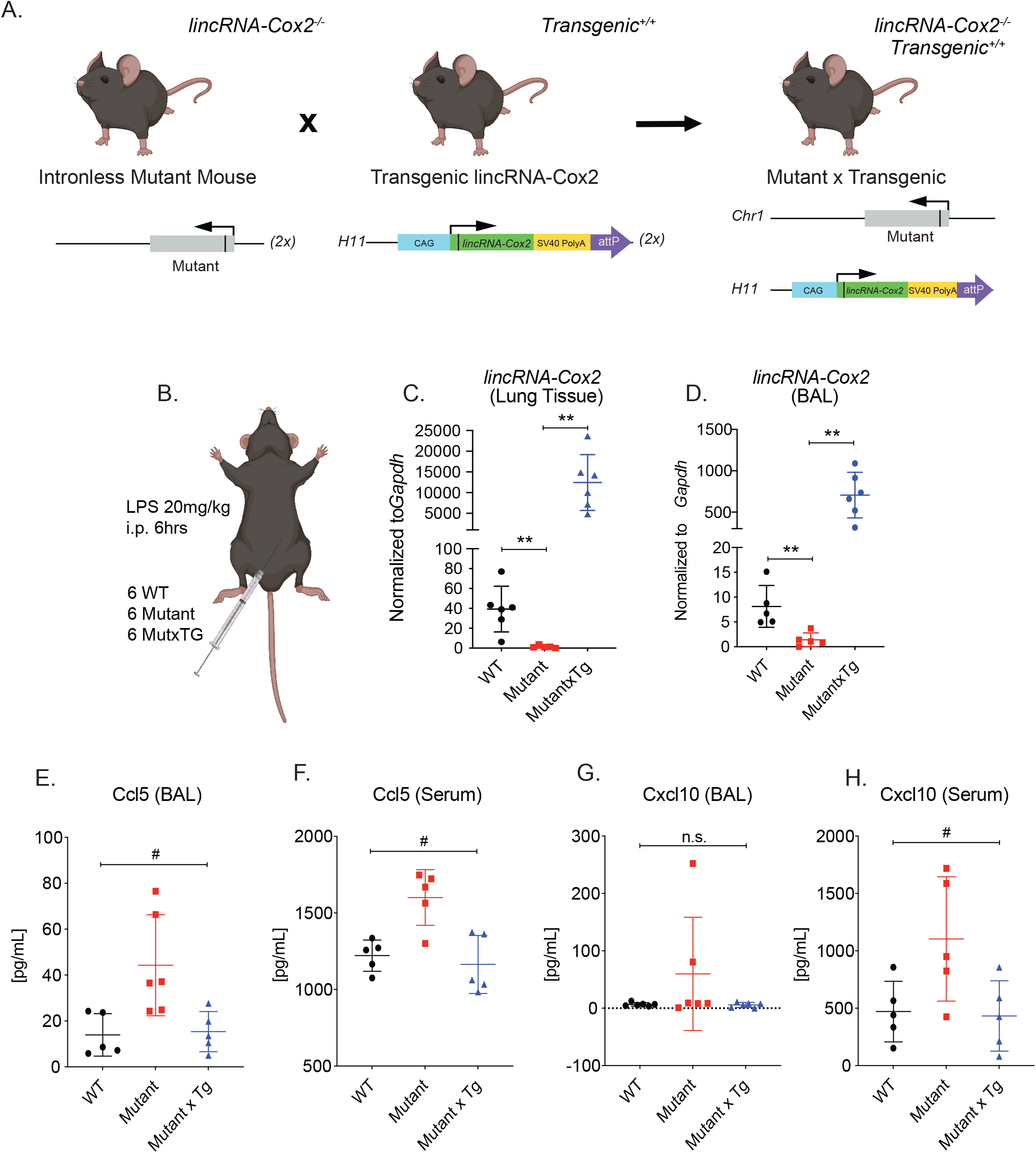
lincRNA-Cox2 functions in *trans* to regulate the innate immune system in a septic shock model. (A) Schematic depicting i.p. route of LPS infection in WT, mutant and transgenic mice. (C) WT, Mutant and TgxMut mice were challenged with 20mg/kg LPS and body temperatures were measured. Mice were sacrificed after 6h, bronchiolar lavage fluid (BAL), lungs and cardiac punctures were performed. BAL and Lungs were harvested for gene expression analysis by RT-qPCR for lincRNA-Cox2 (C-D), Ccl5 (E-F). BAL and isolated serum were sent for multiplex cytokine analysis (G-H). Each dot represents an individual animal. Student’s t-tests were performed using Graphpad Prism7. Asterisks indicate statistically significant differences between mouse lines (*= >0.05, **= >0.01 and *** = >0.005). One-way ANOVA used to determine significance between WT, Mut, and MutxTG mice (#=>0.05).

Next, we wanted to determine if transgenic overexpression of lincRNA-Cox2 can reverse the phenotype observed in the deficient mice during acute lung injury (Figure 5A). We assessed immune cell recruitment in both the BAL and lung tissue by flow cytometry in WT, mutant and MutxTG mice. Again, we found that neutrophils are the only immune cell that is significantly lower in lincRNA-Cox2 mutant mice, while neutrophil recruitment in MutxTG mice return to WT levels (Figure 5B-C). Next, we performed ELISAs on harvested lung tissue, BAL and serum to measure the protein concentration of cytokines and chemokines. We found that Il6, Ccl5, Ccl3, Ccl4, Ccl22 and Ifnb1 are consistently significantly dysregulated in the lincRNA-Cox2 mutant mice and again this phenotype could be rescued back to WT levels by the transgenic overexpression of lincRNA-Cox2 (MutxTG) (Figure 4D-I). Interestingly, Il6, Ccl3, Ccl4 and Ifnb1 are all significantly different in the BAL, while Ccl5 and Ccl22 are significantly dysregulated in the lung tissue only suggesting that lincRNA-Cox2 can impact genes in a cell-specific manner. We find that Tnf is consistently unaffected between all genotypes (Figure 4J). From these data we conclude that lincRNA-Cox2 functions in *trans* to regulate the lung immune response during acute inflammation.

**Figure 5:**
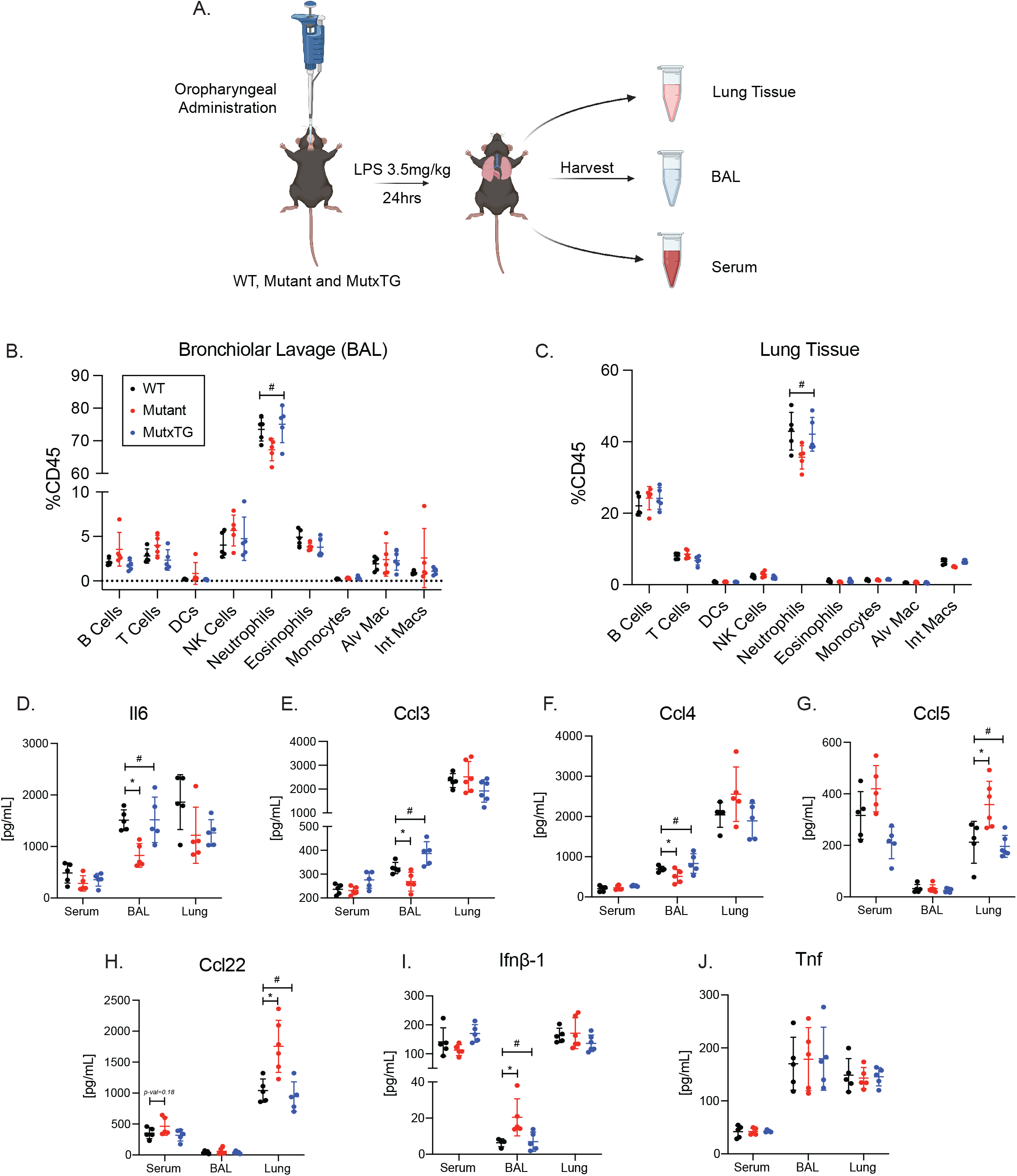
lincRNA-Cox2 regulates the proinflammatory response in the lung in *trans*. (A) Generation of lincRNA-Cox2 MutxTG homozygous mouse. (B) ALI schematic depicting the oropharyngeal route of 3.5mg/kg LPS administration in WT, Mutant, and MutxTG. Mice were sacrificed after 24 h, followed by harvesting lung tissue, serum, and BAL fluid. (C) Body temperatures were measured at 0 h, 3 h, 6 h, and 24 h. (D) BAL and (E) Lung cells were analyzed by flow cytometry to assess recruitment of immune cells in WT and immune cells gated off CD45+. Multiplex cytokine analysis was performed on serum, BAL, and Lung tissue for (F) Il6, (G) Ccl5, (H) Ccl3, (I) Ccl4, (J) Ccl22, (K) Ifnb1 and (L) Tnf. Student’s t-test used to determine significance between WT and Mut mice (*=> 0.05). One-way ANOVA used to determine significance between WT, Mut, and MutxTG mice (#=>0.05).

### LincRNA-Cox2 positively and negatively regulates immune genes in primary alveolar macrophages

In order to understand how lincRNA-Cox2 could be regulating acute inflammation we first wanted to determine which cell types lincRNA-Cox2 is most highly expressed within the lung. First, we utilized publicly available single-cell RNA sequencing (scRNA-seq) data from two LPS-induced lung injury studies (Riemondy *et al*, 2019; Mould *et al*, 2019). Overall, lincRNA-Cox2 was very low in these datasets (SFigure 5A-E). There was a slight increase in expression in all alveolar epithelial type 2 (ATII) cellular populations (SFigure 5D-E). Due to the expression level limitations of publicly available scRNA-seq datasets, we next performed fluorescence activated cell sorting (FACS) to isolate all cell-types of interest in the lung at baseline and following LPS-induced lung injury (Elling *et al*, 2018). Using RT-qPCR we found that lincRNA-Cox2 was most highly expressed in neutrophils at both baseline and following LPS stimulation when normalized to cell count (SFigure 6). Interestingly alveolar macrophages (AMs) were identified as the cell type with the highest induction of lincRNA-Cox2, following LPS stimulation (Figure 6A).

**Figure 6:**
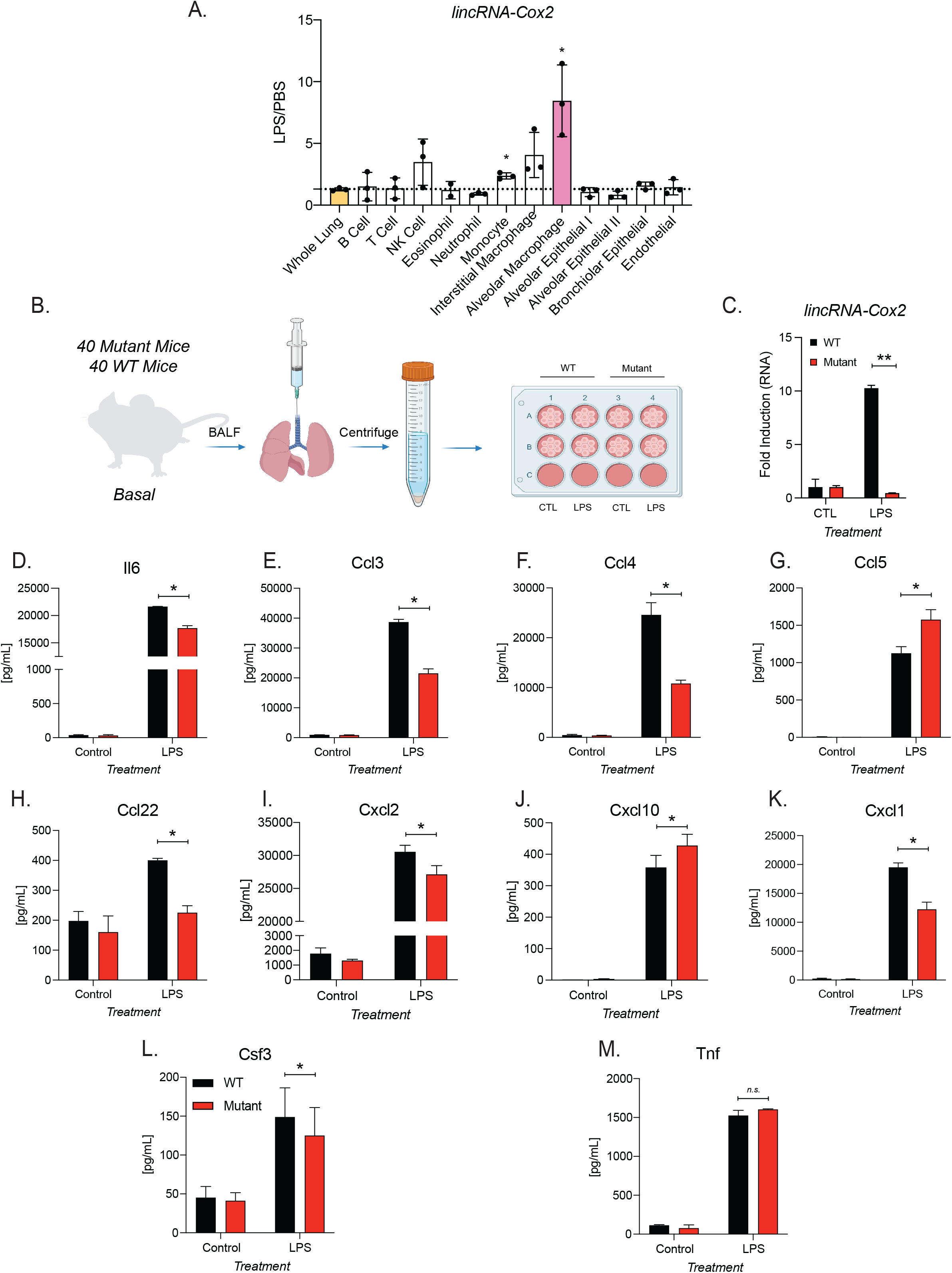
lincRNA-Cox2 is inducible and regulates immune genes both positively and negatively in primary alveolar macrophages. (A) lincRNA-Cox2 was measured in whole lung tissue and several sorted immune and epithelial cells from mice treated with PBS and LPS via an oropharyngeal route. Expression was normalized to PBS. Performed in biological triplicates and student’s t-test was performed between whole lung tissue and each sorted cell. (B) The experimental design is depicted. BALs harvested from 40 WT and 40 Mutant mice, 10 WT or Mutant mice were pooled per well. Cells were treated with LPS for 24 h. (C) lincRNA-Cox2 was measured by RT-qPCR in primary alveolar macrophages. Multiplex cytokine analysis was performed supernatant from primary alveolar macrophages for (D) Il6, (E) Ccl5, (F) Ccl3, (G) Ccl4, (H) Ccl22, (I) Ccl2, (J) Cxcl10 and (K) Cxcl1. Student’s t-test used to determine significance and asterisks indicate statistical significance (*=> 0.05, **>=.01, ***=> 0.0005).

Since lincRNA-Cox2 is most highly induced in AMs we wanted to determine if this was the cell type contributing to the cytokine and chemokine changes in lincRNA-Cox2 deficient (Mut) mice following inflammatory challenge. To do this, we harvested BAL fluid from WT and lincRNA-Cox2 mutant mice to culture primary alveolar macrophages and treated them with LPS for 24 h (Figure 6B). First, lincRNA-Cox2 induction was validated using RT-qPCR with *in vitro* LPS stimulated WT AMs while expression as expected is diminished in the lincRNA-Cox2 deficient AMs (Figure 6B-C). Finally, we assessed the level of cytokine and chemokine expression from primary AMs by ELISA. We confirmed significant dysregulation of Il6, Ccl5, Ccl3, Ccl4 and Ccl22 in primary alveolar macrophages, which are consistent with the *in vivo* data (Figure 6D-H). Additionally, we find novel dysregulation of Cxcl2, Cxcl1, Cxcl10 in alveolar macrophages not detected in our ALI model. Again, Tnf remains consistently unchanged between WT and mutant in AMs and *in vivo* studies (Figure 6M). These data indicate that mechanistically lincRNA-Cox2 is functioning to regulate gene expression primarily within primary alveolar macrophages.

### Peripheral immune cells drive the regulatory role of lincRNA-Cox2 during ALI

From our *in vivo* mouse models, we can conclude that lincRNA-Cox2 functions in *trans* to regulate immune genes and cellular milieu within the lung during ALI. To determine if lincRNA-Cox2 functions through bone-marrow derived immune cells, we performed bone marrow (BM) transplantation experiments utilizing Ubiquitin C (Ubc)-GFP WT bone marrow to enable us to easily track chimerism through measurement of GFP. Ubc-GFP WT BM was transplanted into lincRNA-Cox2 mutant mice and WT mice generating WT→WT and WT→Mut mice (Figure 7A). First, we determined the reconstitution of HSCs by measuring donor chimerism (GFP%) in the peripheral blood (PB) for the duration of the 8 weeks. We find that both the WT→WT and WT→Mut mice have 100% donor reconstitution of granulocytes/ myelomonocytes (GMs) and B cells and ∼75% donor T cells in peripheral blood (Figure 7B-D). While there was a small but significant decrease in T cell reconstitution in WT→Mut mice, we found there is no significant reconstitution difference of GMs or B cells between WT→WT and WT→Mut mice.

**Figure 7:**
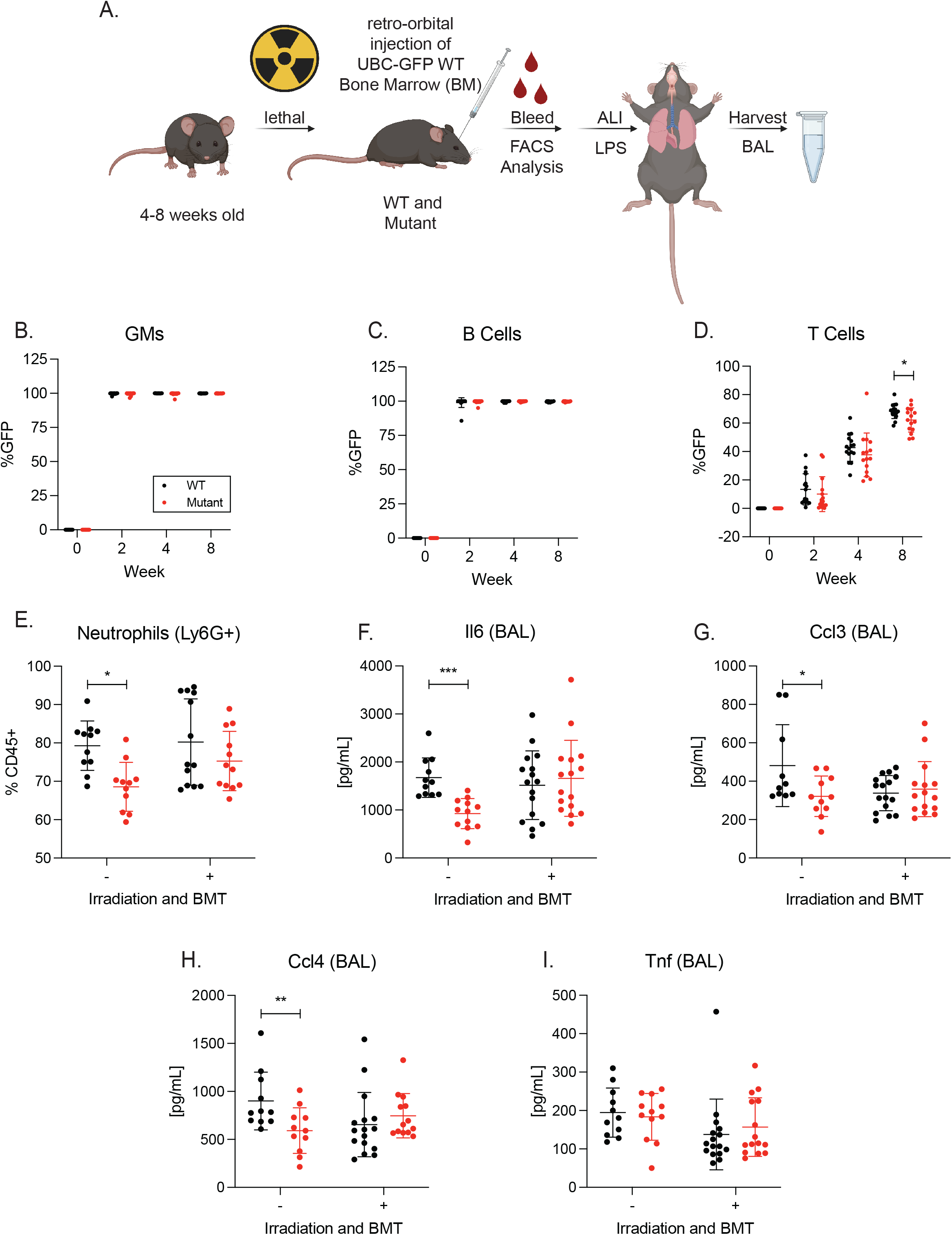
WT Bone marrow transplantation in lincRNA-Cox2 mutant mice rescues the ALI phenotype. (A) WT bone marrow transplantation and ALI experiment schematic. Chimerism was assessed in the peripheral blood by gating for GFP% of (B) granulocytes/myelomonocytes (GMs), (C) B cells and (D) T cells. (E) Percentage of Ly6G+ neutrophil populations were graphed of WT→WT and WT→Mut mice. Multiplex cytokine analysis was performed on BAL for (F) Il6, (G) Ccl3, (H) Ccl4 and (K) Tnf. Data of non-bone marrow transplant (BMT) mice experiments are from Figures 2 and 5. Student’s t-test used to determine significance and asterisks indicate statistical significance (*=> 0.05, **>=.01, ***=> 0.0005).

After 8 weeks when the immune system was fully reconstituted, we performed the LPS ALI model on the chimera WT→WT and WT→Mut mice. We found that the percentage of donor reconstitution within peripheral blood was 100% indicating a successful BM transplantation (Figure 7B-D, SFigure 7) (Hashimoto *et al*, 2013). We found that the decrease in neutrophil recruitment that we identified in the lincRNA-Cox2 mutant mice (Figure 2B, Figure 5B-C, Figure 7E) were rescued in the WT→Mut model back to similar levels to the WT→WT mice (Figure 7E). Furthermore, the altered expression of Il6, Ccl3, and Ccl4 found in the lincRNA-Cox2 mutant mice were also returned to WT levels in the WT→Mut mice (Figure 2C-E, Figure 5D-F, Figure 7F-J). As expected, Tnf acts as a control cytokine showing no difference across genotypes. (Figure 7K). These data suggest that lincRNA-Cox2 is functioning through an immune cell from the bone marrow, most likely alveolar macrophages to regulate acute inflammatory responses within the lung.

## Discussion

LncRNAs are rapidly emerging as critical regulators of biological responses and in recent years there have been several studies showing that these genes play key roles in regulating the immune system (Robinson *et al*, 2020). However very few studies have functionally characterized lncRNAs using mouse models in vivo. We and others have studied the role for lincRNA-Cox2 in the context of macrophages and shown that it can act as both a positive and negative regulator of immune genes (Carpenter *et al*, 2013; Tong *et al*, 2016; Hu *et al*, 2016; Covarrubias *et al*, 2017; Hu *et al*, 2018; Liao *et al*, 2020; Xue *et al*, 2019). We previously characterized two mouse models of lincRNA-Cox2, a knockout (KO) and an intronless splicing mutant (Mut). We identified lincRNA-Cox2 as a *cis* acting regulator of its neighboring protein coding gene Ptgs2 using the KO mouse model. In order to study the role for lincRNA-Cox2 independent of its *cis* role regulating Ptgs2 we generated the splicing mutant (Mut) mouse model. This model enabled us to show that knocking down transcription of lincRNA-Cox2 impacts a number of immune genes including Il6 and Ccl5 in an LPS induced endotoxic shock model. In this current study we make use of the mutant mouse model to study the role for lincRNA-Cox2 in regulating immune responses in the lung, where lincRNA-Cox2 is most highly expressed, both at steady-state and following LPS-induced acute lung injury (ALI). Both neutrophil recruitment and chemokine/cytokine induction are hallmarks of acute lung injury (ALI) (Allen, 2014; Domscheit *et al*, 2020; Ali *et al*, 2020) and here we provide *in vivo* and *in vitro* evidence that lincRNA-Cox2 plays a critical role in these processes.

We find that loss of lincRNA-Cox2 at baseline results in the up- and down-regulation of a number of genes that regulate the immune system and metabolism (Figure 1A-D) indicating that lincRNA-Cox2 is a key transcriptional regulator of gene expression within lung tissue. In addition, we measured immune cells and found that at baseline dendritic cells and B cells were lower in the lincRNA-Cox2 mutant mice. Lung DCs serve as a functional signaling/sensing units to maintain lung homeostasis by orchestrating host responses to benign and harmful foreign substances, while B cells are crucial for antibody production, antigen presentation and cytokine secretion (Wang *et al*, 2019a; Menon *et al*, 2021). Having fewer DC and B cells at steady state could lead to an increased risk of inflammatory diseases (Seys *et al*, 2015; Cook & MacDonald, 2016). This indicates that lincRNA-Cox2 plays an important role in maintaining lung homeostasis since gene expression and cellular abundance are altered by the loss of lincRNA-Cox2.

While we identify lincRNA-Cox2 as a crucial element for maintaining lung homeostasis, we next performed LPS induced acute lung injury (ALI) to assess the importance of lincRNA-Cox2 during active inflammation. Using a 24 h time point, which shows the maximum influx of PMNs and cytokine/chemokine expression (Domscheit *et al*, 2020), we find that loss of lincRNA-Cox2 leads a decrease in neutrophil recruitment and altered cytokine/chemokine expression in both the BAL and serum (Figure 2). In acute lung injury, neutrophils are crucial for bacterial clearance during live infection, repair and tissue remodeling after ALI (Blázquez-Prieto *et al*, 2018; Giacalone *et al*, 2020). Our data suggests that lincRNA-Cox2 plays an important role in neutrophil recruitment and therefore could also play roles in clearance of live bacteria and repair of tissue after resolution of infection. We found that Il6 levels are reduced while Ccl5 levels are increased following ALI in the lincRNA-Cox2 mutant mice. These findings are consistent with our previous *in vitro* and *in vivo* studies (Carpenter *et al*, 2013; Elling *et al*, 2018). Newly, we found that Ccl3 and Ccl4 are positively regulated, while Ccl22 and Ifnb1 are negatively regulated by lincRNA-Cox2 in the lung during ALI. Interestingly, Lee *et al*. reported in humans that the chemokines, CCL3 and CCL4, promote the local influx of neutrophils (Lee *et al*, 2000). Therefore, the decreased expression of Ccl3 and Ccl4 *in vivo* in the lincRNA-Cox2 mutant mice could explain the significant decrease of neutrophil recruitment seen in the BAL (Figure 2B).

To date there remains only a small number of lncRNAs that have been functionally and mechanistically characterized in vivo. In fact, Pnky, Tug1 and Firre are the only other lncRNA studies that show that their phenotype can be rescued in *trans in vivo* (Andersen *et al*, 2019; Lewandowski *et al*, 2019, 2020). From our previous *in vivo* studies, we had concluded that lincRNA-Cox2 functions in *cis* to regulate Ptgs2 in an eRNA manner while it functions in *trans* to regulate genes such as Il6 and Ccl5 (Elling *et al*, 2018). In order to prove that indeed lincRNA-Cox2 can function in *trans* to regulate immune genes we generated a transgenic mouse overexpressing lincRNA-Cox2 from the H11 locus using the TARGATT system (Tasic *et al*, 2011) (Figure 3A). Simply overexpressing lincRNA-Cox2 has minimal impacts on the immune response at baseline or following LPS challenge *in vivo* suggesting that locus specific or inflammatory specific induction of lincRNA-Cox2 is important for its role in regulating immune genes. However, we do note that Il6 levels are higher in the lungs of lincRNA-Cox2 transgenic mice and Csf1 and Il10 are lower in serum following endotoxic shock suggesting that simply overexpressing lincRNA-Cox2 can impact a small number of genes. Our primary goal for generating the transgenic mouse line overexpressing lincRNA-Cox2 was to determine if crossing it to our lincRNA-Cox2 mutant (deficient) mouse generating a MutxTG line (Figure 4A) would rescue the observed phenotypes following LPS challenge using either an intraperitoneal delivery or via oropharyngeal delivery. Interestingly, we find that our MutxTg mice do rescue the phenotype found in both the endotoxic shock model (Figure 4) and LPS induced ALI model (Figure 5), showing definitively that lincRNA-Cox2 regulates gene regulation and cellular recruitment in *trans*. To delve more deeply into exactly how lincRNA-Cox2 is functioning to regulate immune genes in the lung we focused on determining which cell type lincRNA-Cox2 is most highly expressed in. Analysis of scRNA-seq indicated that lincRNA-Cox2 was highly expressed in naive and injured alveolar epithelial type II (AECII) cells (Supplemental Figure 5), however overall lincRNA-Cox2 was difficult to detect in single-cell data probably due to a combination of low expression levels and low read depth. Utilizing FACS and qRT-PCR, we measured the expression of lincRNA-Cox2 in 8 immune cell populations and 4 epithelial/endothelial cell populations and found that lincRNA-Cox2 was most highly expressed in neutrophils (SFigure 6). However, when assessing induction of lincRNA-Cox2 following LPS and normalizing to PBS controls we found it to be most highly expressed in alveolar macrophages with some significant induction also observed in monocytes (Figure 6A). We know alveolar macrophages are critical effector cells in initiating and maintaining pulmonary inflammation, as well as termination and resolution of pulmonary inflammation during acute lung injury (ALI) (Beck-Schimmer *et al*, 2005; Herold *et al*, 2011). Therefore, to determine if the altered gene expression profiles we observed following ALI in the BAL were due to lincRNA-Cox2 expression in alveolar macrophages we cultured primary alveolar macrophages from our WT and lincRNA-Cox2 mutant mice and measured cytokine and chemokine expression (Machiels *et al*, 2017; Nayak *et al*, 2018; Robinson *et al*, 2021). Excitingly, we found decreased expression of Il6, Ccl3 and Ccl4 and increased expression of Ccl5 in the lincRNA-Cox2 mutant alveolar macrophages (Figure 6 D-G). Several other chemokines such as Ccl3, Ccl4, Csf3, Cxcl1 and Cxcl2 were significantly lower in the lincRNA-Cox2 deficient alveolar macrophages and these all are known to play roles in neutrophil influx (Lee *et al*, 2000; Kobayashi, 2008; Metzemaekers *et al*, 2020). These data suggest that lincRNA-Cox2 functions within alveolar macrophages to regulate gene expression including key chemokines that can impact neutrophil infiltration during acute lung injury.

During ALI many of the immune cells that infiltrate the lung including some classes of alveolar macrophages originate from the bone marrow. Therefore, to assess if lincRNA-Cox2 is functioning through bone marrow (BM) derived immune cells we performed BM transplantation chimera experiments in WT and lincRNA-Cox2 mutant mice (Figure 7A). We found that transplanting WT BM into our lincRNA-Cox2 mutant mice completely rescued the neutrophil and cytokine/chemokine phenotype in the BAL (Figure 7B-K). While we aimed to determine if lincRNA-Cox2 functions either through resident (SiglecF+) or recruited (Siglec F-) alveolar macrophages, we found that both populations were composed of >70% donor BM (SFigure 8I-J) indicating that radiation obliterated resident SiglecF+ alveolar cells which become repopulated with donor cells from the bone marrow (Guilliams *et al*, 2013; Hashimoto *et al*, 2013; Misharin *et al*, 2017; Collins *et al*, 2020; Gangwar *et al*, 2020). These experiments enable us to conclude that lincRNA-Cox2 expression originating from the bone marrow can function to control immune responses in the lung, since BM derived immune cells transplanted into lincRNA-Cox2 mutant mice are able to rescue the phenotype driven by loss of lincRNA-Cox2 in the lung.

In conclusion, in this study we show, through multiple mouse models, that lincRNA-Cox2 is functioning in *trans* in alveolar macrophages to regulate immune responses within the lung. This study provides an additional layer of mechanistic understanding highlighting that lncRNAs can contribute to the delicate balance between maintenance of homeostasis and induction of transient inflammation within the lung microenvironment.

**Table 1:**
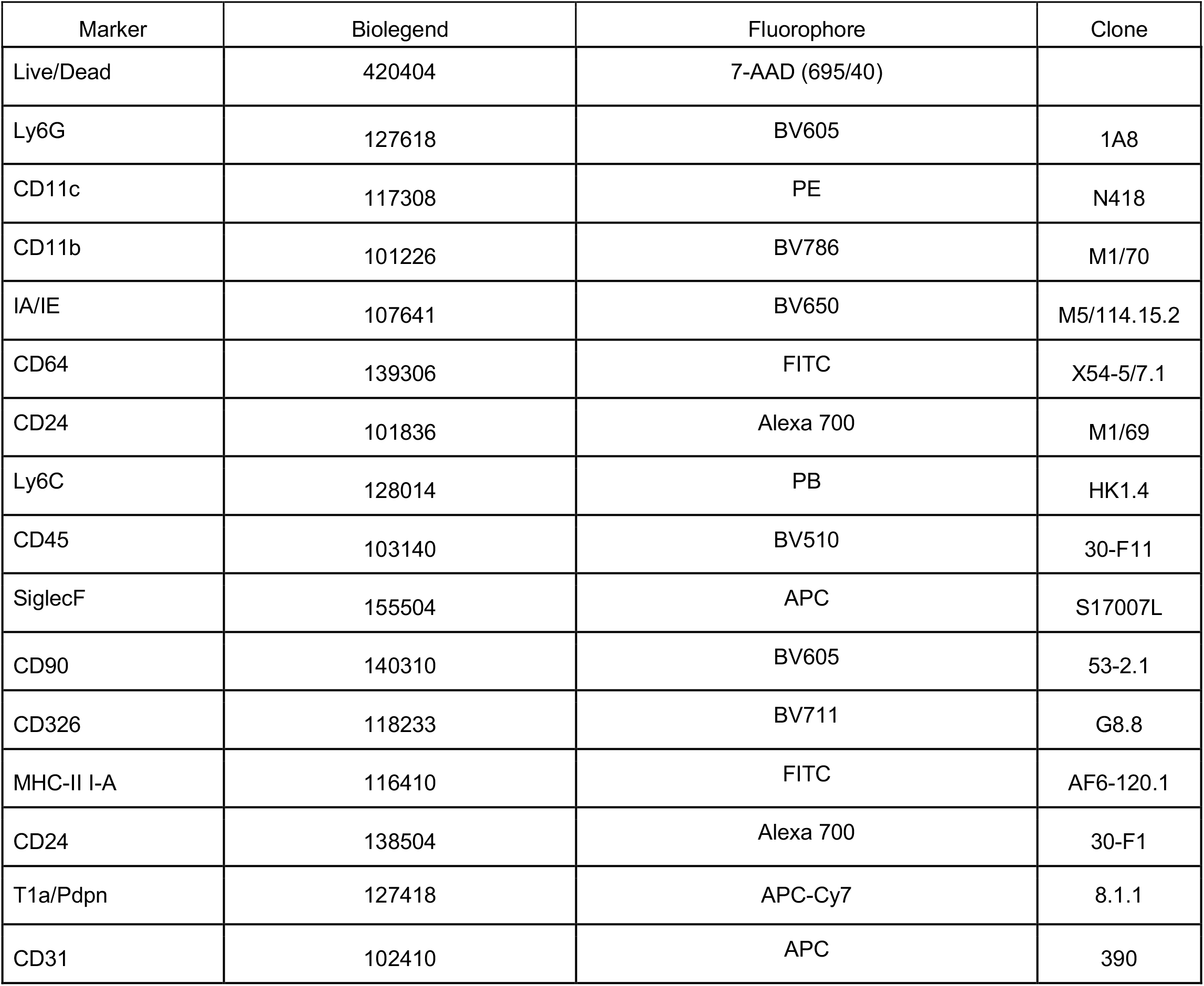
Antibodies used in all Non-Chimera Experiments.

**Table 2:** Antibodies used in Chimera Experiments.

| Marker | Biolegend | Fluorophore | Clone |
| --- | --- | --- | --- |
| Live/Dead | 420404 | 7-AAD (695/40) |  |
| Ly6G | 127612 | PB | 1A8 |
| CD11c | 117308 | PE | N418 |
| CD11b | 101226 | BV786 | M1/70 |
| IA/IE | 107628 | APC-Cy7 | M5/114.15.2 |
| CD64 | 139314 | PE-Cy7 | X54-5/7.1 |
| CD24 | 101827 | BV605 | M1/69 |
| SiglecF | 155504 | APC | S17007L |
| CD45 | 103128 | A700 | 30-F11 |

## Acknowledgments

We thank the UCSC Flow Cytometry Core Facility, RRID:SCR_021149, for technical assistance. Additionally, we thank Biorender for creating a platform to easily generate figures using “Biorender.com”.

## Author Contribution

E.K.R. designed, performed, and analyzed all molecular biology and *in vivo* experiments for all figures. A.K.W. and D.M.P. assisted with flow cytometry and sorting experiments, as well as helped design and perform chimera transplantation experiments. BS set-up and performed experiments for Figure 3. M.M.S. and H.H. assisted with all *in vivo* experiments. B.M. analyzed bulk RNA-sequencing experiments. V.S. helped with analyzing RT-qPCR experiments in Figure 5. L.S. analyzed all scRNA-sequencing datasets. E.C.F. helped with the design of BM transplantation and chimera experiments. E.K.R. and S.C. conceived and coordinated the project. E.K.R. and S.C. wrote the manuscript with input from all other co-authors.

## Methods

### Mice

Wild-type (WT) C57BL/6 mice were purchased from the Jackson Laboratory (Bar Harbor, ME) and bred at the University of California, Santa Cruz (UCSC). All mouse strains, including lincRNA-Cox2 mutant (Mut), transgenic (Tg) and MutxTg mice, were maintained under specific pathogen-free conditions in the animal facilities of UCSC and protocols performed in accordance with the guidelines set forth by UCSC and the Institutional Animal Care and Use Committee.

### Generation of lincRNA-Cox2 Transgenic (Tg) and MutxTg Mice

lincRNA-Cox2 transgenic mice were generated by using a site-specific integrase-mediated approach described previously (Tasic *et al*, 2011). In brief, TARGATT mice in the C57/B6 background contain a CAGG promoter within the Hipp11 (H11) locus expressing the full length lincRNA-Cox2 (variant 1) as previously cloned (Carpenter *et al*, 2013) generated at the Gladstone (UCSF). These mice were then genotyped using the same TARGATT approach of PCR7/8 (PR432:GATATCCTTACGGAATACCACTTGCCACCTATCACC, SH176:TGGAGGAGGACAAACTGGTCAC, SH178:TTCCCTTTCTGCTTCATCTTGC). The lincRNA-Cox2 transgenic mice were then crossed with the lincRNA-Cox2 mutant mice and bred to homozygosity to generate MutxTg mice. For genotyping to assess homozygosity of Mutant we used the primer sets of MutF: ATGCCCAGAGACAAAAAGGA and MutR: GATGGCTGGATTCCTTTGAA, as well as the 3-primer set stated above.

### ALI model

Age- and sex matched WT, lincRNA-Cox2 mutant and lincRNA-Cox2 MutxTg mice were treated with 3.5mg/kg using the orophyrangeal intratracheal administration technique. The model of LPS insult via oropharyngeal administration into the lung was previously described in detail (Allen, 2014; Nielsen *et al*, 2018; Ehrentraut *et al*, 2019). Briefly, mice were sedated using an isoflurane chamber (3% for induction, 1-2% for maintenance), then 60-75ul of 3.5mg/kg of LPS (from strain O111:B4) or PBS (control) were administered using a pipette intratracheally. 24 h after LPS treatment, mice were sacrificed using CO2 and serum, BALF and lung were harvested for either cellular assessment by flow cytometry, RNA expression or sent to EVE technologies for cytokine/chemokine protein analysis.

### LPS Shock model

Age- and sex matched wild-type, lincRNA-Cox2 mutant mice, lincRNA-Cox2 Tg and lincRNA-Cox2 MutxTg (Elling *et al*, 2018) (8-12 weeks of age) were injected i.p. with 20 mg/kg LPS (O111:B4). For gene expression analysis and cytokine analysis, mice were euthanized 6 h post injection. Blood was taken immediately postmortem by cardiac puncture. Statistics were performed using GraphPad prism.

### Transplantation reconstitution Assays

Reconstitution assays were performed, as previously stated by Poscablo *et al*. (Poscablo *et al*, 2021), by transplanting double-sorted HSCs (200 per recipient) from Ubc-GFP+ whole BM and transplanting into congenic C57BL/6 WT and lincRNA-Cox2 deficient mice via retro-orbital intravenous transplant. We also transplanted double-sorted MkPs (22,000 per recipient) from C57Bl6 into Ubc-GFP+ hosts. Hosts were preconditioned with sub-lethal radiation (∼750 rads) using a Faxitron CP160 X-ray instrument (Precision Instruments).

### Harvesting Bronchiolar Lavage Fluid (BAL)

Bronchoalveolar Lavage Fluid (BALF) was harvested as previously stated by Cloonan *et al*. (Cloonan *et al*, 2016). 40 mice were euthanized by CO2 narcosis, the tracheas cannulated, and the lungs lavaged with 0.5-ml increments of ice-cold PBS eight times (4 ml total), samples were combined in 50 ml conical tubes. BALF was centrifuged at 500 *g* for 5 min. 1 ml red blood cell lysis buffer (Sigma-Aldrich) was added to the cell pellet and left on ice for 5 min followed by centrifugation at 500 *g* for 5 min. The cell pellet was resuspended in 500 μl PBS, and leukocytes were counted using a hemocytometer. Specifically, 10 μl was removed for cell counting (performed in triplicate) using a hemocytometer. Cells were plated in sterile 12 well plates at 5e5/well (total of 8 wells) and use complete DMEM with 25 ng/ml supGM-CSF.

### Lung Tissue Harvesting for cytokine measurement

Mice were humanely sacrificed, and their lungs were excised. The whole lungs were snap frozen and homogenized, and the resulting homogenates were incubated on ice for 30 min and then centrifuged at 300 × *g* for 20 min. The supernatants were harvested, passed through a 0.45-μm-pore-size filter, and used immediately or stored at −70°C, then sent to EVE for measurements of cytokines/chemokines.

### Cell culture of Primary AMs

24 h post-BALF isolation, media was removed and fresh complete DMEM with 25 ng/ml supGM-CSF is added (Robinson *et al*, 2021). All cells that adhere to the surface of the plate are considered alveolar macrophages (AM) as previously determined by Chen *et al*. (Chen *et al*, 1988). After new media is added, AMs are stimulated with 200 ng/ml LPS (Sigma, L2630-10MG). Harvest supernatant 6 h post-stimulation. Harvested supernatant was sent to Eve technologies for cytokine analysis. Statistics were performed using GraphPad prism.

### RNA isolation, cDNA synthesis and RT-qPCR

Total RNA was purified from cells or tissues using Direct-zol RNA MiniPrep Kit (Zymo Research, R2072) and TRIzol reagent (Ambion, T9424) according to the manufacturer’s instructions. RNA was quantified and assessed for purity using a nanodrop spectrometer (Thermo Fisher). Equal amounts of RNA (500 to 1,000 ng) were reverse transcribed using iScript Reverse Transcription Supermix (Bio-Rad, 1708841), followed by qPCR using iQ SYBR Green Supermix reagent (Bio-Rad, 1725122) with the following parameters: 50 °C for 2 min and 95 °C for 2 min, followed by 40 cycles of 95 °C for 15 s, 60 °C for 30 s, and 72 °C for 45 s, followed by melt-curve analysis to control for nonspecific PCR amplifications. Oligos used in qPCR analysis were designed using Primer3 Input version 0.4.0 (https://bioinfo.ut.ee/primer3-0.4.0/).

Gene expression levels were normalized to Gapdh or Hprt as housekeeping genes.

*Primers Used:*

Gapdh F- CCAATGTGTCCGTCGTGGATC

Gapdh R - GTTGAAGTCGCAGGAGACAAC

HPRT F - TGCTCGAGATGTCATGAAGG

HPRT R - ATGTCCCCCGTTGACTGAT

lincRNACox2 F - AAGGAAGCTTGGCGTTGTGA

lincRNACox2 R – GAGAGGTGAGGAGTCTTATG

### ELISA

The concentration of Il6 and Ccl5 levels in the serum and BAL of WT, lincRNA-Cox2 mutant mice, lincRNA-Cox2 Tg and lincRNA-Cox2 MutxTg mice were determined using the DuoSet ELISA kits (R&D, DY1829 and DY478) according to the manufacturer’s instructions.

### Lung Tissue Harvesting for Cellular Analysis

Mice were humanely sacrificed, and their lungs were excised. Lung was inflated with a digestion solution containing 1.5mg/ml of Collagenase A (Roche) and 0.4mg/ml DNaseI (Roche) in HBSS plus 5% fetal bovine serum and 10mM HEPES. Trachea was tied off with 2.0 sutures. The heart and mediastinal tissues were carefully removed, and the lung parenchyma placed in 5ml of digestion solution and incubated at 37°C for 30 minutes with gently vortexing every 8–10 minutes. Upon completion of digestion, 25ml of PBS was added; and the samples were vortexed at maximal speed for 30 seconds. The resulting cell suspensions were strained through a 70um cell strainer and treated with ACK RBC lysis solution. Then the cells were stained using the previously published immune (Yu *et al*, 2016) and epithelial (Nakano *et al*, 2018) cellular panels.

### Flow Cytometry Analysis and Sorting

After cells were isolated and counted, ∼2×10^6 cells per sample were incubated in blocking solution containing 5% normal mouse serum, 5% normal rat serum, and 1% FcBlock (eBiosciences, San Diego, CA) in PBS and then stained with a standard panel of immunophenotyping antibodies (See Supplemental Table for a list of antibodies, clones, fluorochromes, and manufacturers) for 30 minutes at room temperature (Yu *et al*, 2016). Data was acquired and compensation was performed on the BD Aria II and Attune NxT (Thermo Fisher) flow cytometer at the beginning of each experiment. Data was analyzed using Flowjo v10. Cell sorting was performed on a BD Aria II. The collected cells were harvested for RNA and RT-qPCR was performed to measure lincRNA-Cox2. Analysis was performed using FlowJo analysis software (BD Biosciences).

### RNA-sequencing Analysis

Generation of the RNAseq data (GSE117379) and analysis of differential gene expression has been described previously (Elling *et al*, 2018). RNA-seq 50bp reads were aligned to the mouse genome (assembly GRCm38/mm10) using TopHat. The Gencode M13 gtf was used as the input annotation. Differential gene expression specific analyses were conducted with the DESeq2 R package. Specifically, DESeq2 was used to normalize gene counts, calculate fold change in gene expression, estimate p-values and adjusted p-values for change in gene expression values, and to perform a variance stabilized transformation on read counts to make them amenable to plotting.

### single cell RNA-sequencing Analysis

Generation of the sc-RNAseq data (Gene Expression Omnibus accession number GSE120000 and GSE113049) and analysis of gene expression has been described previously (Mould *et al*, 2019; Riemondy *et al*, 2019).

#### Plotting Expression across cells

For overlaying expression onto the tSNE plot for single genes or for the average across a single panel of genes, we plotted normalized expression along a continuous color scale, with the extreme color values being set to the 5th and 95th quantile expression values. The bubble plot and heatmap show scaled normalized expressions along a continuous color scale. We produced the heatmap using Heatmap3 (Zhao *et al*, 2014).

---

**Supplemental Figure 1:**
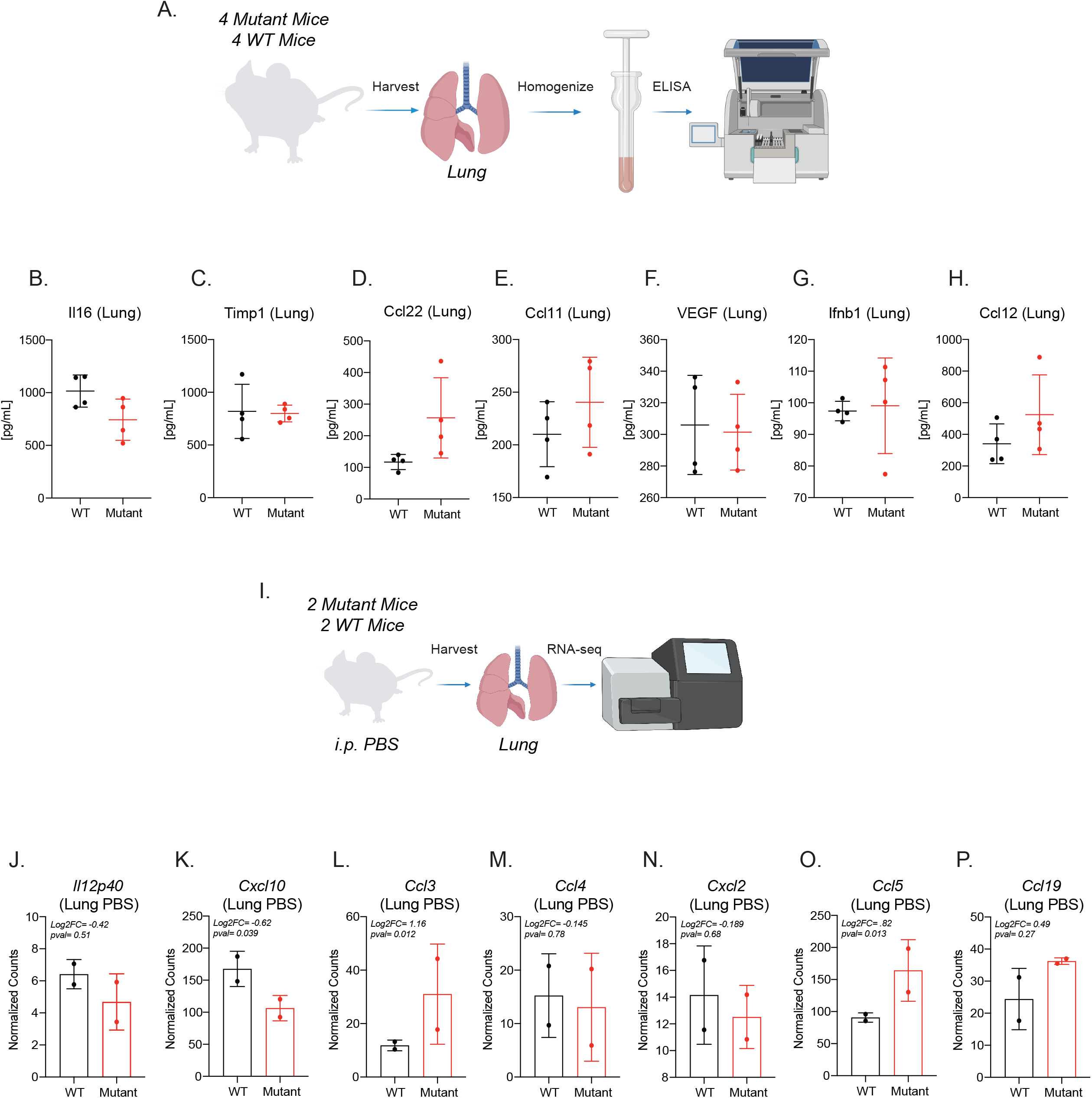
Characterization of immune pathways in lincRNA-Cox2 mutant at baseline. (A) Schematic of cytokine analysis of lung homogenates from WT and mutant mice. Multiplex cytokine analysis was performed on lung homogenates for (B) Il16, (C) Timp1, (D) Ccl22, (E) Ccl11, (F) Vegf, (G) Ifnb1 and (H) Ccl12. (I) Schematic of RNA-seq analysis of WT and Mutant lungs at baseline. Normalized counts for (J) Il12p40, (K) Cxcl10, (L) Ccl3, (M) Ccl4, (N) Cxcl2, (O) Ccl5 and (P) Ccl19. Student’s t-test used to determine significance and asterisks indicate statistical significance (*=> 0.05, **>=.01, ***=> 0.0005).

**Supplemental Figure 2:**
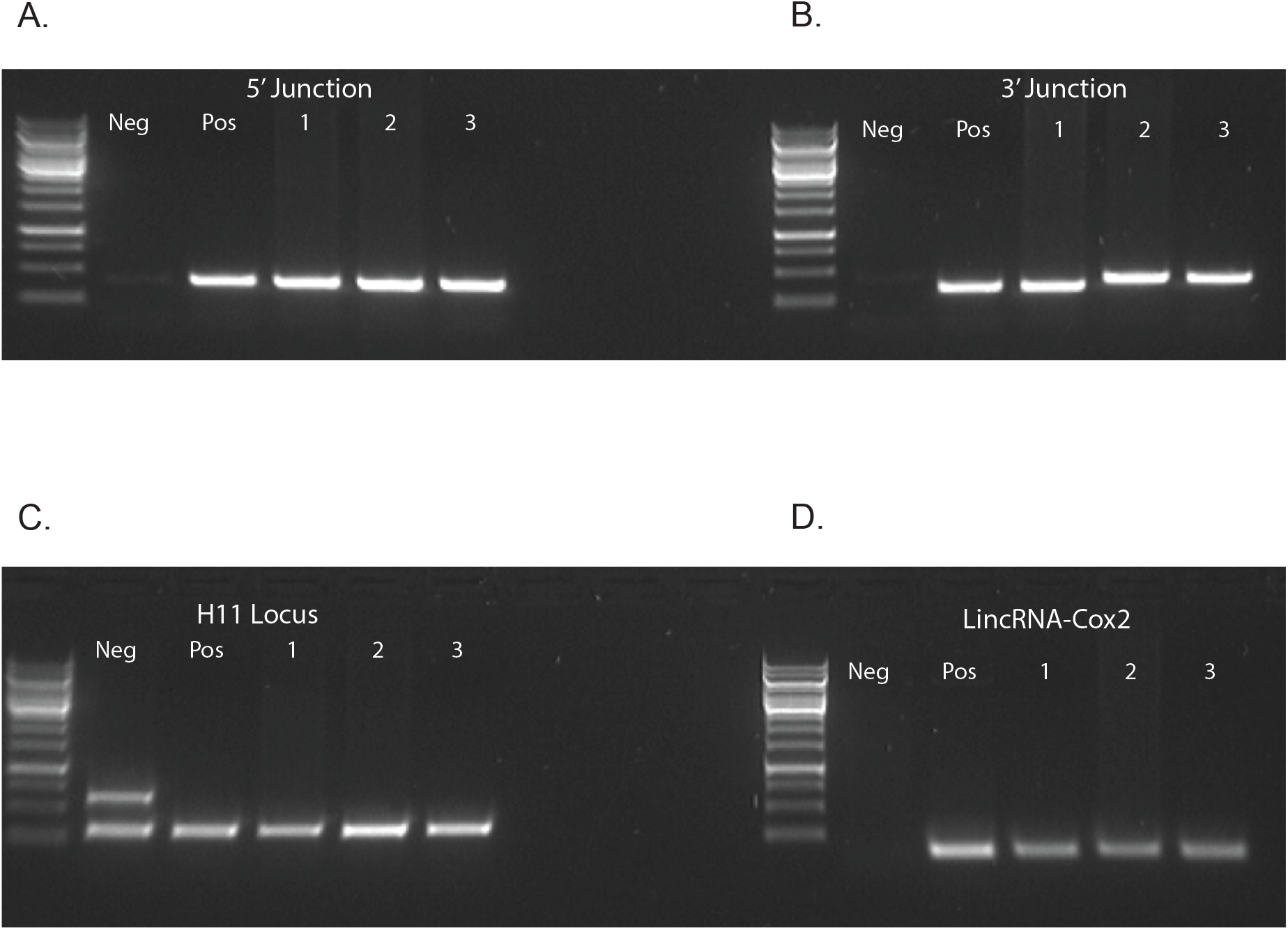
PCR strategy of transgenic lincRNA-Cox2 mouse using the TARGATT system. Generation of transgenic mouse with the Targatt system. This process allowed recombination at the H11 locus. Specific primer sets were used to confirm correct integration of lincRNA-Cox2. The strategy used to generate the lincRNA-Cox2 transgenic mice was adapted from Tasica *et al,* PNAS 2011 and genotyping (A-D) strategy was utilized to confirm homozygosity.

**Supplemental Figure 3:**
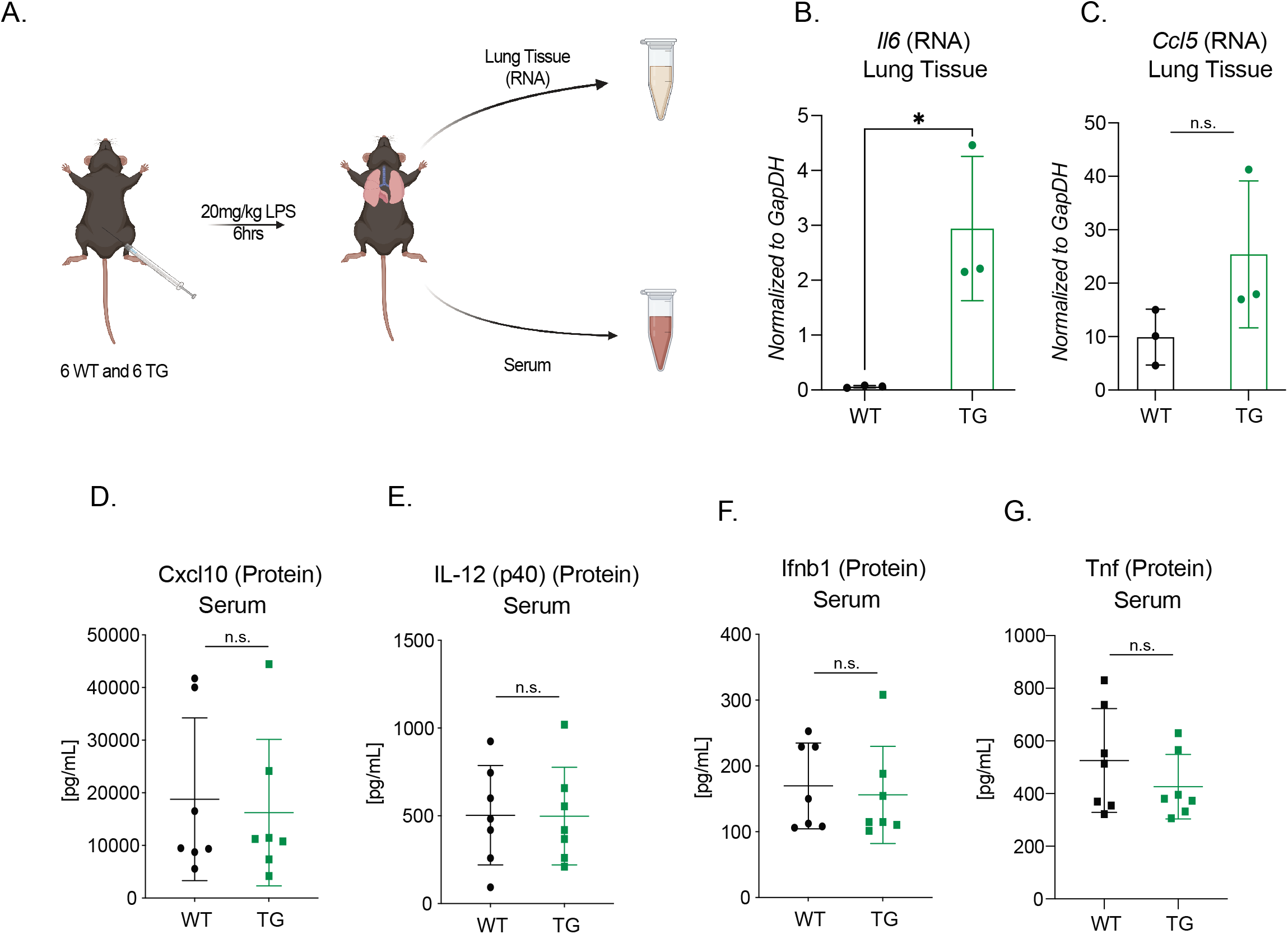
Overexpression of lincRNA-Cox2 *in vivo* does not regulate acute inflammation. (A) Schematic of WT and Transgenic septic shock model. RNA was isolated from lung tissue to measure mRNA expression of (B) Il6 and (C) Ccl5, normalized to GapDH. Serum was harvested to measure (D) Cxcl10, (E) Il12p40, (F) Ifnb1 and (G) Tnf by ELISA. Student’s t-test used to determine significance and asterisks indicate statistical significance (*=> 0.05, **>=.01, ***=> 0.0005).

**Supplemental Figure 4:**
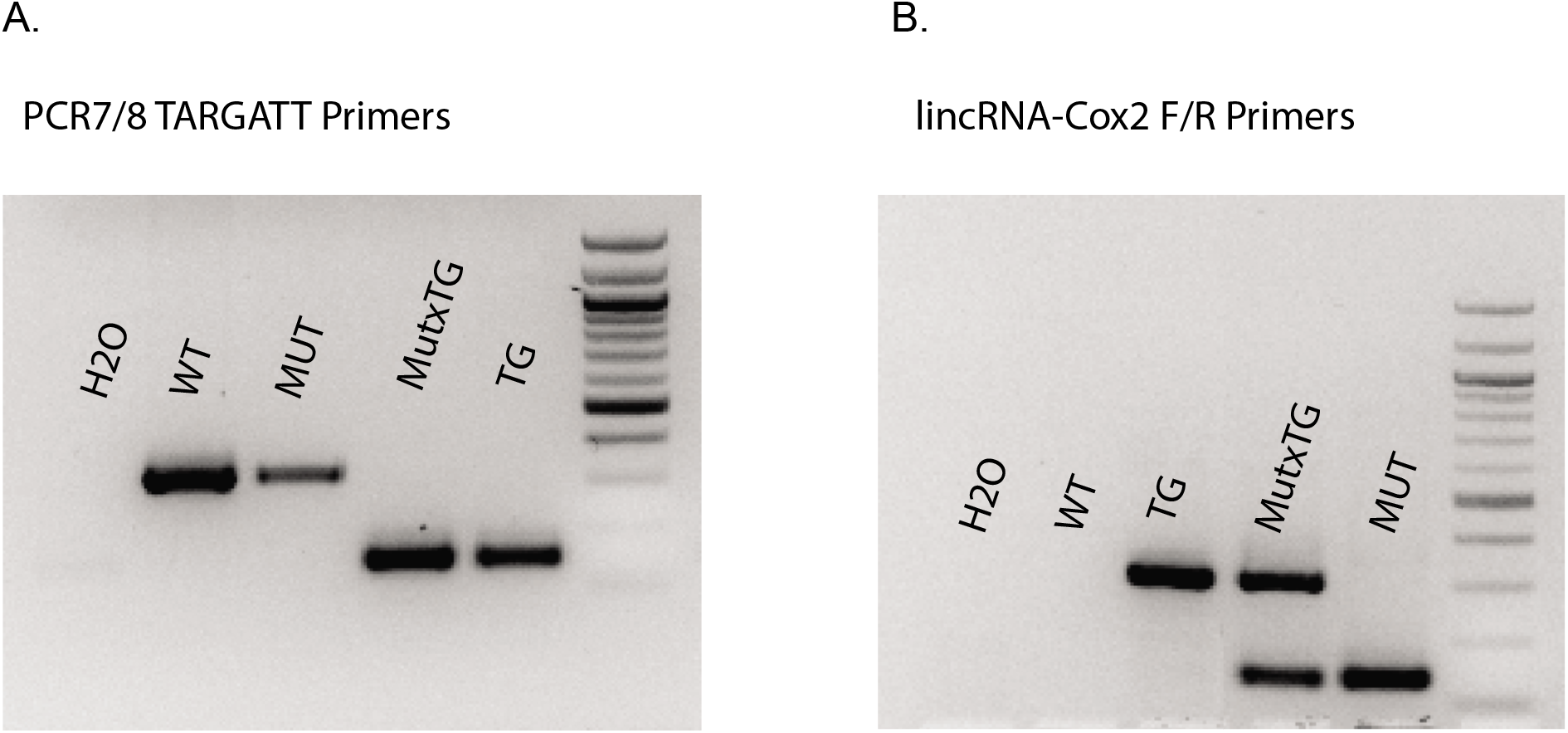
Genotyping strategy for MutxTG lincRNA-Cox2 mouse. (A) PCR7/8 TARGATT primers were utilized to check for homozygosity. (B) lincRNA-Cox2 specific primers were utilized to assess homozygosity of mutant mouse allele.

**Supplemental Figure 5:**
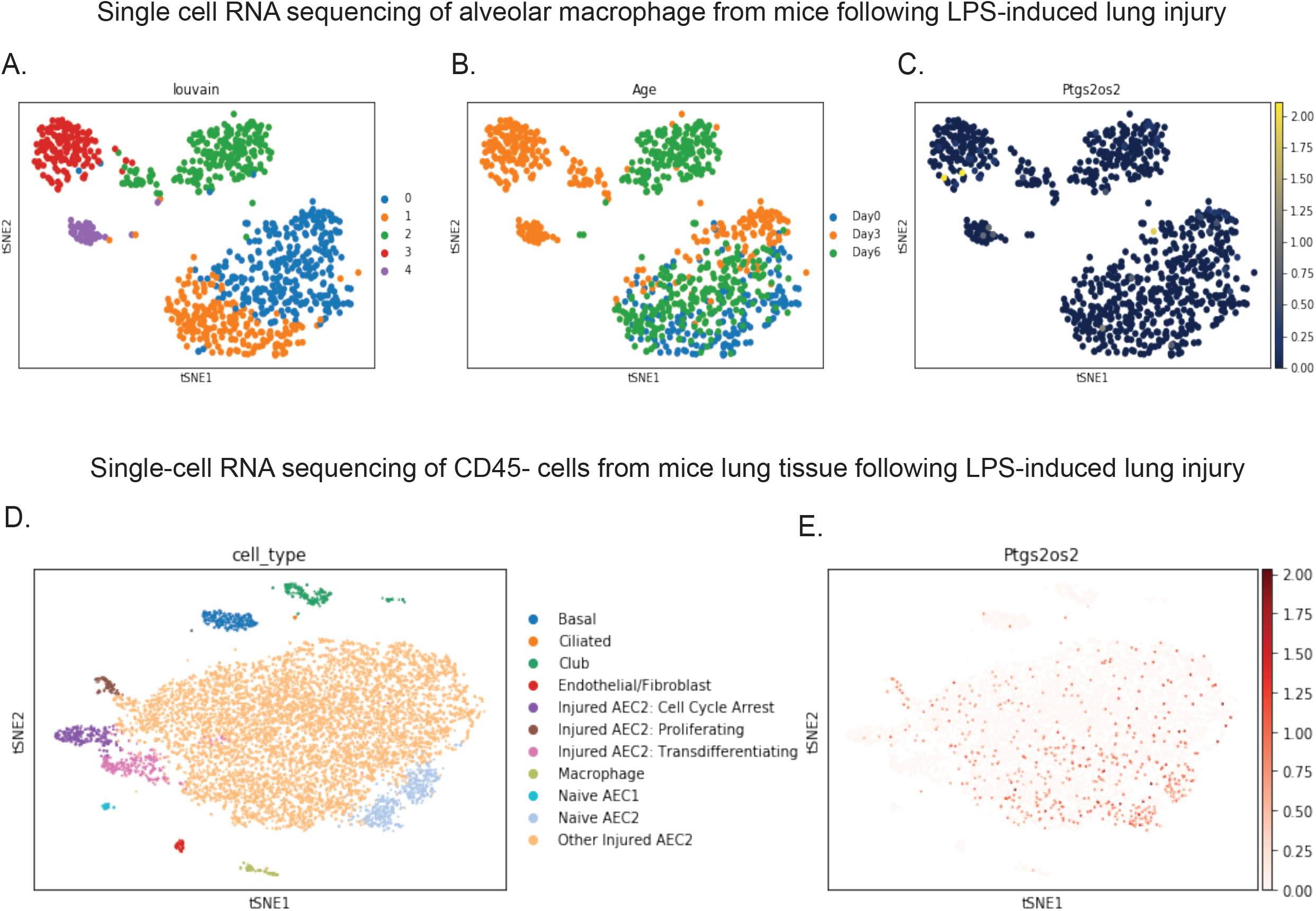
scRNA-seq does not significantly identify cell-type specific lincRNA-Cox2 expression in the lung during acute inflammation. Mould *et al*. generated scRNA-seq of alveolar macrophages from WT mice pre- and post-LPS induced acute lung injury. (A) tSNE plots were generated indicating 5 distinct populations. Then tSNE plots were utilized to examine (B) day of LPS stimulation and (C) lincRNA-Cox2 (ptgs2os2) expression. Riemondy *et al*. generated scRNA-seq of all CD45-cells from WT mice pre- and post-LPS induced acute lung injury. (D) tSNE plots were generated and clusters were colored based on cell type and (E) lincRNA-Cox2 (ptgs2os2) expression.

**Supplemental Figure 6:**
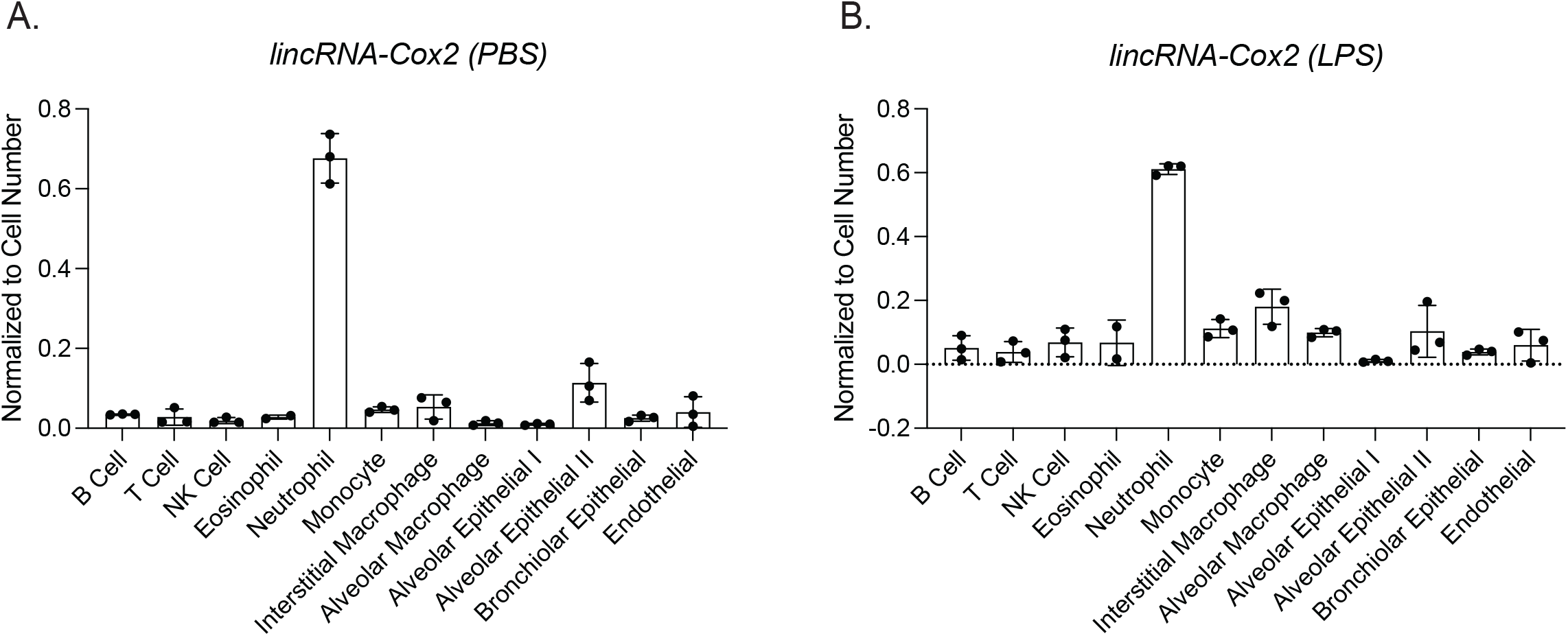
lincRNA-Cox2 expression in sorted cells from PBS or LPS treated mice. lincRNA-Cox2 was measured in whole lung tissue and a number of sorted immune and epithelial cells from mice treated with (A) PBS and (B) LPS via an oropharyngeal route. Performed in biological triplicates and student’s t-test was performed between whole lung tissue and each sorted cell.

**Supplemental Figure 7:**
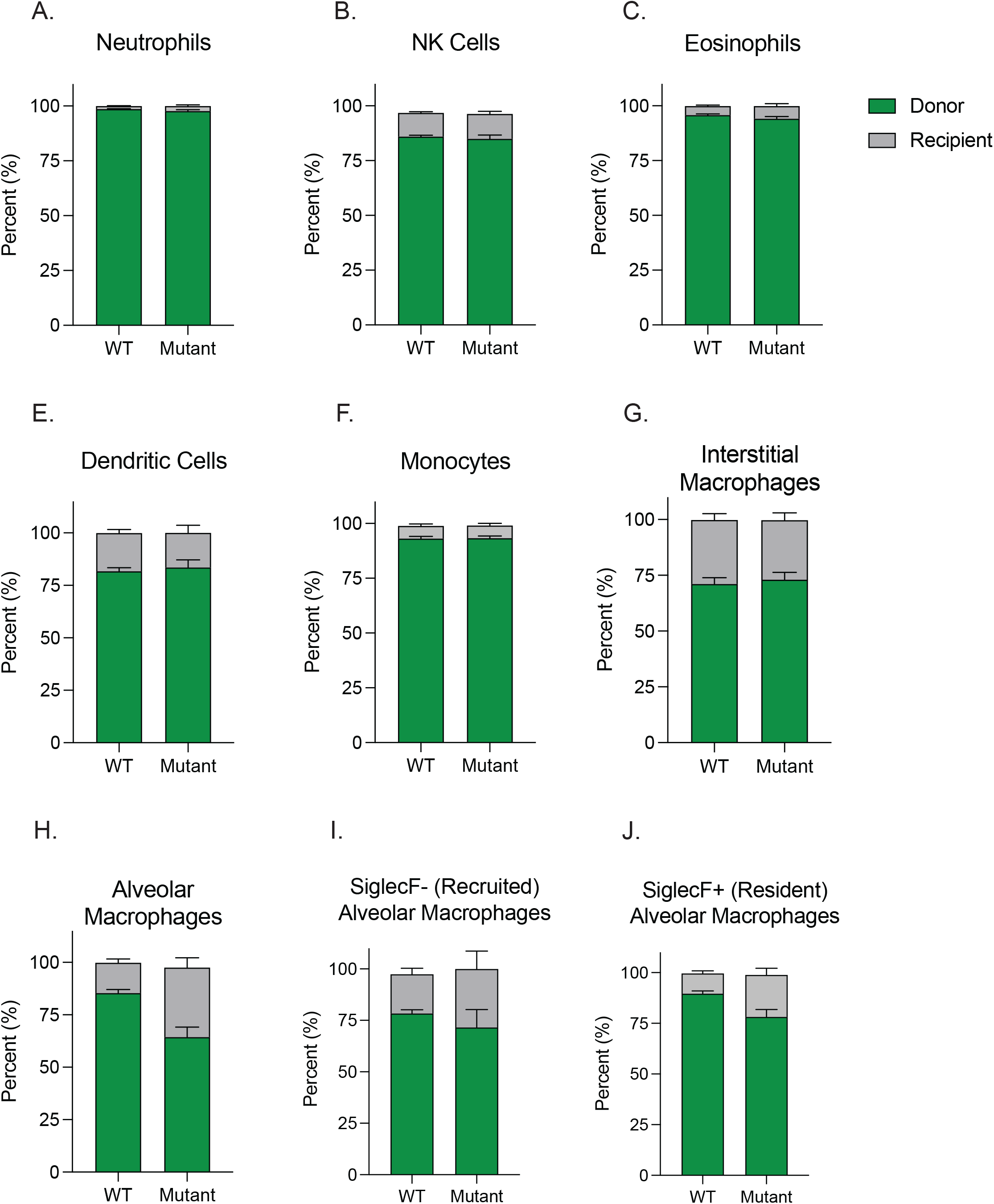
Reconstitution of donor BM in the BAL of WT and Mutant mice. Donor (green) and recipient (grey) percentage were assessed for (A) Neutrophils, (B) NK cells, (C) Eosinophils, (D) Dendritic cells, (E) Monocytes, (F) Eosinophils, (G) Interstitial macrophages, (H) Alveolar macrophages, (I) Recruited alveolar macrophages and (J) Resident Alveolar macrophages.

